# *In vitro* live cell imaging reveals nuclear dynamics and role of the cytoskeleton during asymmetric division of pollen mitosis I in *Nicotiana benthamiana*

**DOI:** 10.1101/2025.03.25.644957

**Authors:** Yoko Mizuta, Masako Igarashi, Tomomi Shinagawa, Ikuma Kaneshiro, Daisuke Kurihara

**Affiliations:** Institute of Transformative Bio-Molecules (WPI-ITbM), Nagoya University, Furo-cho, Chikusa-ku, Nagoya, Aichi 464-8601, Japan; Institute for Advanced Research (IAR), Nagoya University, Furo-cho, Chikusa-ku, Nagoya, Aichi 464-8601, Japan; Department of Physics, Graduate School of Science, Nagoya University, Furo-cho, Chikusa-ku, Nagoya, Aichi 464-8602, Japan; Research Center for Computational Science, Institute for Molecular Science, National Institutes of Natural Sciences, 38 Nishigo-Naka, Myodaiji, Okazaki 444-8585, Japan

**Keywords:** Pollen, Biolistics, Cell culture, Confocal microscopy, Actin, Tubulin, *Nicotiana benthamiana*

## Abstract

Pollen is a male gametophyte of angiosperms. Following meiosis, the microspore undergoes an asymmetric division called pollen mitosis I (PMI), which produces two cells of different sizes: a large vegetative cell and a small generative cell. Polarized nuclear migration and positioning during PMI are important for successful pollen development and cell differentiation. However, analyzing the pollen development process in real-time is challenging in many model plants with tricellular pollen, including *Arabidopsis* and rice. In this study, we established a method for live confocal imaging of microtubule and actin dynamics using suspension cultures with biolistic delivery of plasmid DNAs during PMI in *Nicotiana benthamiana* (Bentham’s tobacco), containing bicellular pollen. Pharmacological studies have indicated that actin filaments are crucial for microspore nuclear positioning before PMI, cell plate expansion during cytokinesis, and chromatin dispersion in vegetative cell nucleus after PMI. By contrast, inhibition of microtubule assembly resulted in abnormal chromosome segregation and nuclear behavior after PMI, although nuclear positioning and asymmetric division were observed. Our *in vitro* live cell imaging system for PMI provides insights into the importance of cytoskeletal regulation in asymmetric division and differentiation during pollen development.

## Introduction

A multicellular organism comprises multiple cells with different functions and roles that are created through cell division and subsequent cell differentiation. Asymmetric division is a fundamental mechanism that generates cellular diversity in animals and plants. Pollen, the male gametophyte of angiosperms, is produced through asymmetric division. During microgametogenesis, a single-cell microspore undergoes an asymmetric division called pollen mitosis I (PMI), which produces two unequal daughter cells (Hafidh & Honys, 2021). These large and small daughter cells differentiate into vegetative and generative cells with completely different cellular identities and biological functions (Figure 1A). Only one generative cell undergoes pollen mitosis II (PMII), which results in the formation of two sperm cells. Asymmetric division during PMI is essential for the differentiation of generative cells and sexual reproduction in angiosperms. Polarized nuclear migration and nuclear positioning along the polar axis are necessary for PMI, which is precisely controlled in asymmetrically dividing cells (Twell et al., 1998). In *Arabidopsis* microspores, the polar axis is defined during the tetrad stage (Erdtman, 1952). Asymmetric division along the polar axis is achieved in three sequential steps: (1) the microspore nucleus migrates toward the wall pole (future generative pole) during interphase, (2) the mitotic spindle assembles near the nucleus and then orients perpendicular to the microspore wall along the polar axis during metaphase, and (3) finally, in the telophase, a smaller generative cell is formed at the generative pole, whereas a larger vegetative cell is formed at the inner pole (vegetative pole). Morphological, pharmacological, and genetic studies suggest that cytoskeletal elements are essential in this process (Brown & Lemmon, 2000). Microtubules and actin filaments are important for polarized nuclear migration and the maintenance of their position in the microspore (Van Lammeren et al., 1985; Van Lammeren et al., 1989). Microtubule inhibitors or centrifugation can induce aberrant symmetric division in the PMI, resulting in the formation of two equal-sized vegetative cells without generative cells (Tanaka & Ito, 1981; Terasaka & Niitsu, 1987; Twell et al., 2002). *Arabidopsis gemini pollen* (*gem*) mutants exhibit symmetric and partial divisions of the PMI (Park et al., 1998; Park et al., 2004). GEM3, which encodes the conserved AUGMIN subunit 6 that is involved in microtubule nucleation, was isolated from these mutants (Oh et al., 2016). Microtubules and actin filaments are considered important for PMI (Van Lammeren et al., 1989). Actin microfilaments have been identified in the spindle and phragmoplast regions of *Brassica napus* (Gervais et al., 1994) and are also implicated in maintaining the position of the microspore nucleus in *Nicotiana tabacum* (Zonia et al., 1999). These cytoskeletal elements, including PMI, are crucial in the plant cell cycle. However, studies that have examined these dynamics during the PMI of living microspores are limited (Daghma et al., 2014). Common model plants such as *Arabidopsis* and rice contain tricellular pollen (Williams et al., 2014). Tricellular pollen completes PMII by anthesis, indicating that PMI is completed at an earlier developmental stage. This is one of the reasons why pollen culture in these model plants is difficult. By contrast, generating transformants from non-model plants using bicellular pollen is difficult and time-consuming. Therefore, traditional methods such as dye staining and immunostaining using fixed pollen have been used to study microgametogenesis (Van Lammeren et al., 1985). To analyze the dynamics of the cytoskeleton during PMI, a live analysis method and rapid study of gene function using bicellular pollen are required.

**Figure 1.**
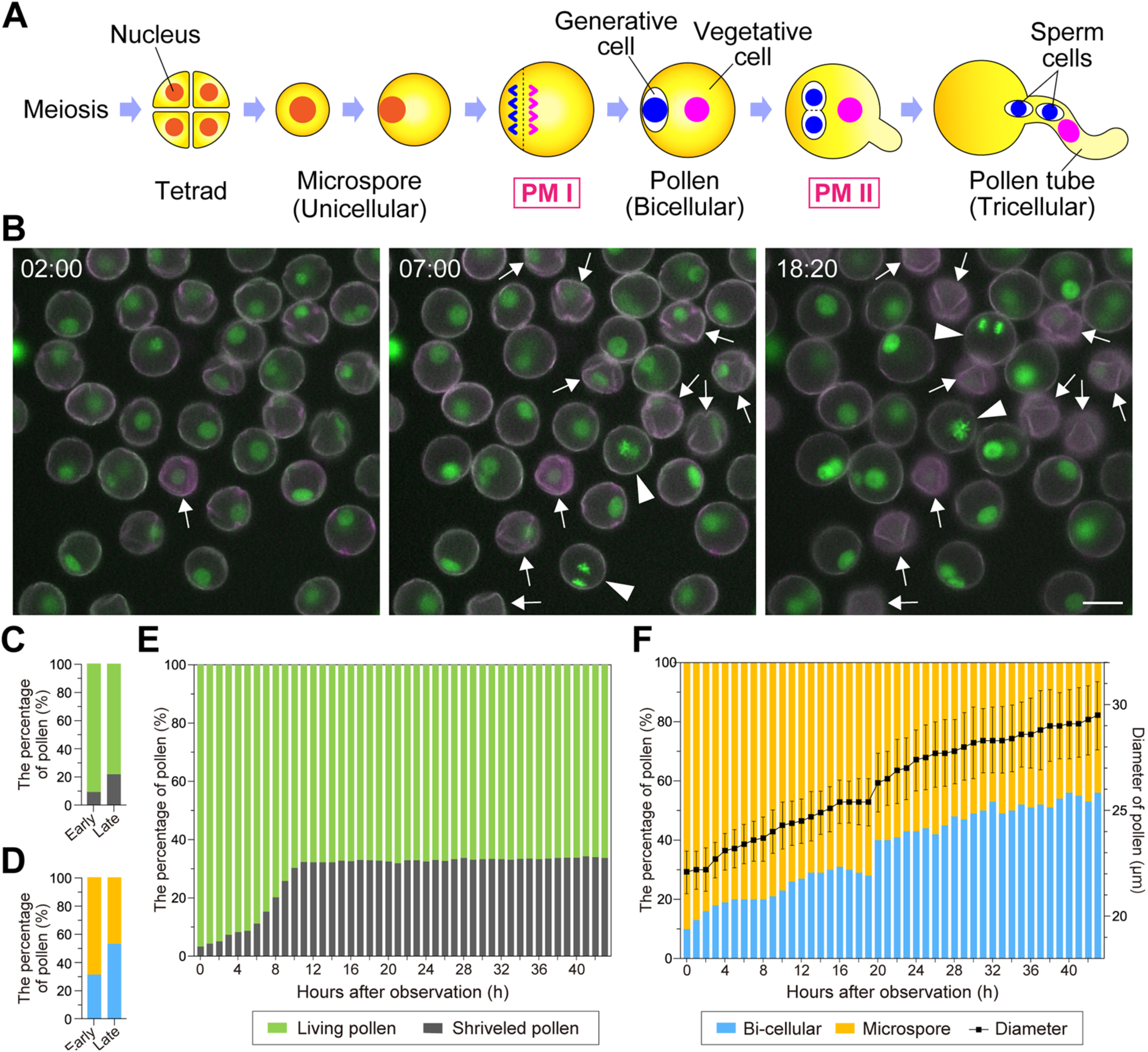
Pollen development and pollen mitosis I of *Nicotiana benthamiana in vitro*. (A) Schematic representation of pollen development in *N. benthamiana*. The pollen mother cell is divided by meiosis to produce haploid pollen tetrads and subsequent microspores. A highly asymmetric division, termed pollen mitosis I (PMI), results in a large vegetative cell and small generative cell as two unequal daughter cells. A plasma membrane surrounds the generative cell. The pollen generative cell is engulfed by the vegetative cell, forming a “cell within a cell” structure. After pollen germination, a generative cell divides into two sperm cells via symmetric division, which is termed pollen mitosis II (PMII). (B) *In vitro* live imaging of PMI in *N. benthamiana*. Nuclei were labeled with *AtUBQ10p::H2B-mClover* (green). The pollen autofluorescence is shown in magenta. Arrowheads indicate pollen from the metaphase to telophase in PMI. Arrows indicate pollen with red autofluorescence and a shrunken shape, which was defined as developmental arrest. The numbers stamped in each image indicate the time (hh:mm) from the beginning of the observation. Scale bar, 20 μm. (C) Pollen viability at each hour *in vitro* was measured by time-lapse imaging. Green bar, living pollen labeled with green fluorescent protein; gray bar, shrunken pollen labeled with red autofluorescence. (D) Developmental stage and diameter of pollen from *AtUBQ10p::H2B-mClover* after each hour of culture. The line graph shows the mean diameter of the 10 pollen grains showing PMI, and the error bars indicate standard deviations. Yellow bar, bicellular pollen; blue bar, microspores. (E, F) Pollen viability (E) and development (F) at each stage *in vivo,* without culture. A total of 877 mid-stage pollen grains and 1,131 late-stage pollen grains were measured. The meanings of the colors are the same as in (C) and (D). Early refers to early-stage pollen (4–5 days before anthesis); Late refers to late-stage pollen (2–3 days before anthesis).

In this study, we established a method to study cytoskeletal dynamics during PMI using *Nicotiana benthamiana* (Bentham’s tobacco) microspores based on the suspension culture method. *N. benthamiana,* which contains bicellular pollen (Russell & Jones, 2015), is the most extensively used experimental host in plant virology for the analysis of plant–pathogen interactions (Goodin et al., 2008). We observed that *N. benthamiana* microspores undergo PMI and develop into mature pollen under *in vitro* conditions, similar to the related species, *N. tabacum* (Tupý et al., 1991). Transiently introducing genes into the pollen via particle bombardment is also feasible, as previously reported (Kaneshiro et al., 2022; Nagahara et al., 2021). Marker plasmids were transiently introduced into the microspores to visualize the cytoskeleton, and microtubule and actin localization were analyzed before and after PMI. The effects of the inhibitors on nuclear migration, polarization, and cytokinesis were examined using imaging. This study elucidates the regulatory mechanisms underlying cytoskeletal dynamics, asymmetric division, and subsequent differentiation during pollen development.

## Results

### Pollen viability and cytokinesis during PMI in *N. benthamiana*

We developed a live imaging method to observe nuclear dynamics and the cytoskeleton during PMI in *N. benthamiana* (Figure 1A). The developmental stage of pollen is not synchronized with stages 1 and 2 (Steinbachová et al., 2021). Therefore, the pollen population in a single anther contains both microspores and immature bicellular pollen. Live imaging of the microspores showed that some microspores developed into bicellular pollen through PMI (Movie 1), whereas others exhibited a loss of nuclear fluorescence signals and a shrunken shape, which appeared to arrest their development (Figure 1B, arrows). Pollen viability, pollen developmental stage, and cell growth were analyzed to investigate the effects of pollen culture. The analysis of 502 early-stage and 542 late-stage microspores showed that the aborted pollen were 9.4% and 21.8%, respectively (Figure 1C). Pollen viability was examined at each hour after incubation using live imaging. The examination of 404 to 607 pollen grains per hour revealed that the aborted pollen initially accounted for 3.3%, subsequently increasing to 32.2% at 12 h from the beginning of observation (Figure 1E). This ratio was slightly higher than that observed without incubation (Figure 1C). The aborted pollen remained constant at approximately 32%–34% from 11 to 43 h following culture, indicating that some factors determine cell viability during the first 11 h of culture (Figure 1E).

Next, the number of nuclei in the viable pollen was measured to investigate the timing of PMI under culture conditions. When 429 early-stage and 473 late-stage microspores were examined from anthers in the bud, the bicellular pollen were 31.2% and 53.1%, respectively (Early and Late in Figure 1D). The bicellular pollen was 10.0% at the beginning of the observation period and then gradually increased to 56.3% at 43 h after culture (Figure 1F). This gradual increase indicates that each microspore developed into bicellular pollen through PMI without synchrony, even under *in vitro* culture conditions similar to *in vivo* conditions (Figure 1D). The pollen diameter was also analyzed using 10 randomly selected bicellular pollen grains that developed during culture. The average diameter was 22.1 ± 1.0 μm at the beginning of the observation, which increased linearly to 29.5 ± 1.6 μm after 43 h of culture, indicating that pollen cells developed during the culture period (Figure 1F).

### Live imaging of microtubule and actin dynamics in PMI using biolistic delivery

During PMI, the establishment of cell polarity and subsequent asymmetric division occurs in two steps: first, the microspore nucleus migrates toward the microspore wall during future division, and second, the mitotic spindle assembles to ensure that the spindle axis is established perpendicular to the microspore wall for asymmetric division (Twell et al., 1998). Cytoskeletal elements play fundamental roles in nuclear positioning and migration (Yi & Goshima, 2022). To analyze the cytoskeletal structure of the microspores during PMI, transient visualization of the nucleus, microtubules, and actin filaments was performed using biolistic delivery of plasmid DNA (Nagahara et al., 2021). *UBQ10p::H2B-tdTomato* plasmid DNA was introduced into microspores from the nuclear marker line. The pollen suspension from 1.5 anthers containing 9,117 pollen grains was poured into a glass-bottom dish (Figure 2A). After 20 h of incubation, 221 of these pollen grains had nuclei labeled with tdTomato (Figure 2A and 2B). Therefore, the efficiency of plasmid delivery into *N. benthamiana* microspores was 2.42%. Some of the introduced pollen grains exhibited bicellular pollen characteristics, indicating that the microspores introduced with plasmid DNA using biolistic delivery were capable of PMI under *in vitro* conditions.

**Figure 2.**
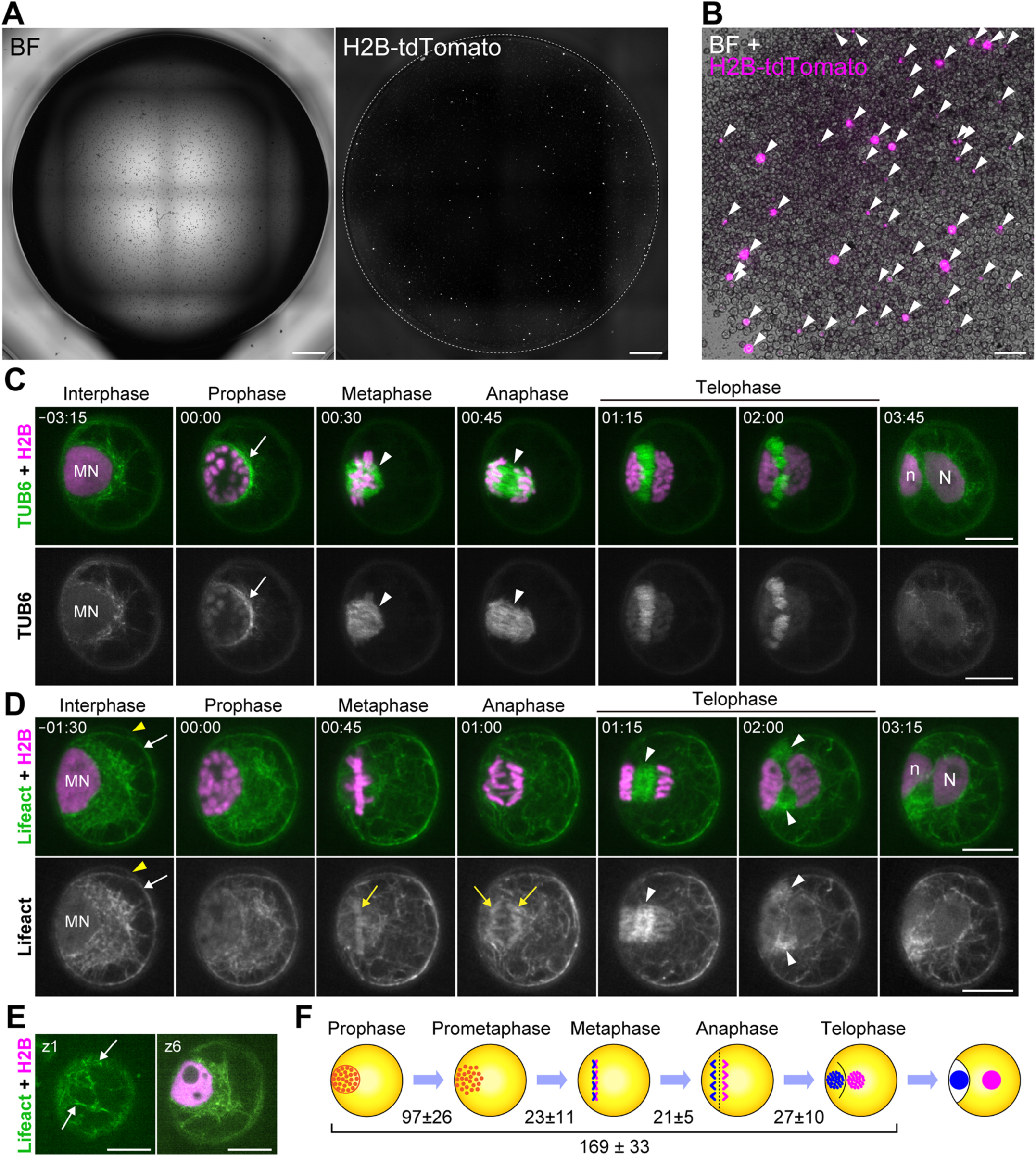
Confocal live imaging of cytoskeletal elements during PMI. (A) Brightfield and fluorescence images of the pollen after bombardment. Microspores from three anthers of wild-type *N. benthamiana* were used for bombardment with the plasmid DNA *UBQ10p::H2B-tdTomato*, and the pollen suspension was then divided into two wells of a glass-bottom dish. Images show one of these wells after 20 h of incubation. (B) Merged brightfield and fluorescence images of bombarded pollen after 20 h of incubation. (C, D) Confocal live imaging of microtubule and actin dynamics during PMI by transiently introducing plasmid markers. The *xy*-maximum projection images in 1-μm steps with two planes are shown. The time at which the prophase began was defined as 0 min. Asterisks, white arrows, and white arrowheads indicate microspore nuclei, peripheral localization of microtubules on the nucleus, and appearance of microtubules on the spindle, respectively. Yellow arrowheads and arrows in (D) indicate autofluorescence of the cell wall and localization of actin on chromosomes, respectively. (E) Cortical actin networks in pollen plasma membranes. The first and sixth z-planes are presented. (F) Time course of each phase of the PMI. Microspore nuclei, vegetative cells, and generative cells are shown in orange, magenta, and blue, respectively. The time was measured for 14 pollen grains in confocal live imaging captured at 15 min intervals. N, vegetative cell nucleus; n, generative nucleus. Scale bars: 1 mm (A), 100 μm (B), and 10 μm (C–E).

Similarly, we transiently introduced a mixture of two plasmid DNAs: *UBQ10p::H2B-tdTomato*, and a microtubule marker containing the *Arabidopsis TUB6* gene (Sasaki et al., 2023). Confocal live imaging revealed the localization dynamics of the microtubules during PMI (Movie 2). During interphase, microtubule filaments were observed in the cytoplasm and localized to the periphery of the microspore nucleus, which became more evident in the subsequent prophase (arrow in Figure 2C; Prophase). Most of these cytoplasmic signals disappeared and formed spindles after nuclear envelope breakdown, and the spindles aligned and separated the chromosomes during metaphase and anaphase (arrowheads in Figure 2C; Metaphase and Anaphase). During telophase, the phragmoplast formed from the remnants of the anaphase spindle and expanded centrifugally to separate the cells (Figure. 2C; Telophase). Microtubules were dispersed in the cytoplasm following the completion of cytokinesis.

We also transiently introduced a mixture of two plasmid DNAs: *UBQ10p::H2B-tdTomato,* and the actin marker *UBQ10pro:Lifeact-mGFP* (Kijima et al., 2025). Actin filaments were similarly localized at the periphery of the microspore nucleus (Movie 3) but were more extensively distributed in the cytoplasm than in the microtubules (Figure 2D; Interphase). In contrast to the microtubules, cortical actin networks were also observed in the plasma membrane (white arrows in Figure 2D and 2E; Interphase), which were separated from the cell wall in the microspore (yellow arrowheads in Figure 2D). During prophase, weak signals were detected on the chromosome until anaphase (yellow arrows in Figure 2D; Metaphase and Anaphase). Strong signals were then localized to the phragmoplast during telophase (white arrowheads in Figure 2D; Telophase), which dispersed into the cytoplasm. Based on these live observations, we measured the time course of each phase in the PMI using 14 time-lapse images taken at 20-min intervals under *in vitro* conditions. The beginning of prophase was defined as 0 min, and the average duration of prophase, prometaphase, metaphase, and anaphase were 97 ± 26, 23 ± 11, 21 ± 5, and 27 ± 10 min, respectively (Figure 2F).

### Disruption of actin filaments impaired the maintenance of polar positioning of the nucleus before PMI, resulting in the failure of asymmetric division

Colchicine inhibits microtubule polymerization resulting in the symmetric division of tobacco microspores during PMI and produces two equally sized daughter cells (Eady et al., 1995). Nuclear positioning at the generative pole is maintained by an actin network (Zonia et al., 1999). To investigate whether microtubules and actin filaments regulate nuclear migration and asymmetric division, microspores isolated from the nuclear marker line were cultured in the presence of the microtubule formation inhibitor, colchicine, or the actin polymerization inhibitor, latrunculin B (LatB). In the control group, the nucleus was located at the center of the microspore and migrated to the generative pole (Movie 4 and Figure 3A; Control). When colchicine was added to the medium at a final concentration of 0.001% (w/v), polar migration of the microspore nucleus to the generative pole was observed in many pollen grains, and some pollen grains exhibited disruption of chromosome organization during metaphase (Movie 4 and Figure 3A; Colchicine). To investigate the role of actin filaments in nuclear migration before PMI, we disrupted them in LatB microspores. In the control microspores, the nucleus was located at the center at an early stage and then migrated to the generative pole (Movie 4 and Figure 3A; Control). When LatB was added to the medium at a final concentration of 25 μM, the positioning of the microspore nucleus was impaired, resulting in division at the center of the microspore. As shown in Movie 4 and Figure 3A, the nucleus migrated to the generative pole, but its position was not maintained, and then floated to the center of the microspore (Figure. 3A; LatB, 00:00). Subsequently, division occurred at a position different from that of the generative pole, resulting in the formation of a cell plate at the center of the pollen, which also impaired cell plate expansion and cytokinesis (arrows in Figure 3A; 25 μM LatB). Time-lapse images were analyzed to investigate the distance between the cell division site and the cell center during anaphase. When the cell division site was measured from the center of the microspore, nuclear migration toward the generative pole and subsequent asymmetric division were observed in control microspores (Movie 5). Similar nuclear migration and division occurred in colchicine-treated microspores, whereas division occurred at shorter distances, that is, at more cell centers in LatB-treated microspores (Figure 3B). This indicates that actin filaments, but not microtubules, are required to maintain the nucleus positioned at the generative pole in the microspore.

**Figure 3.**
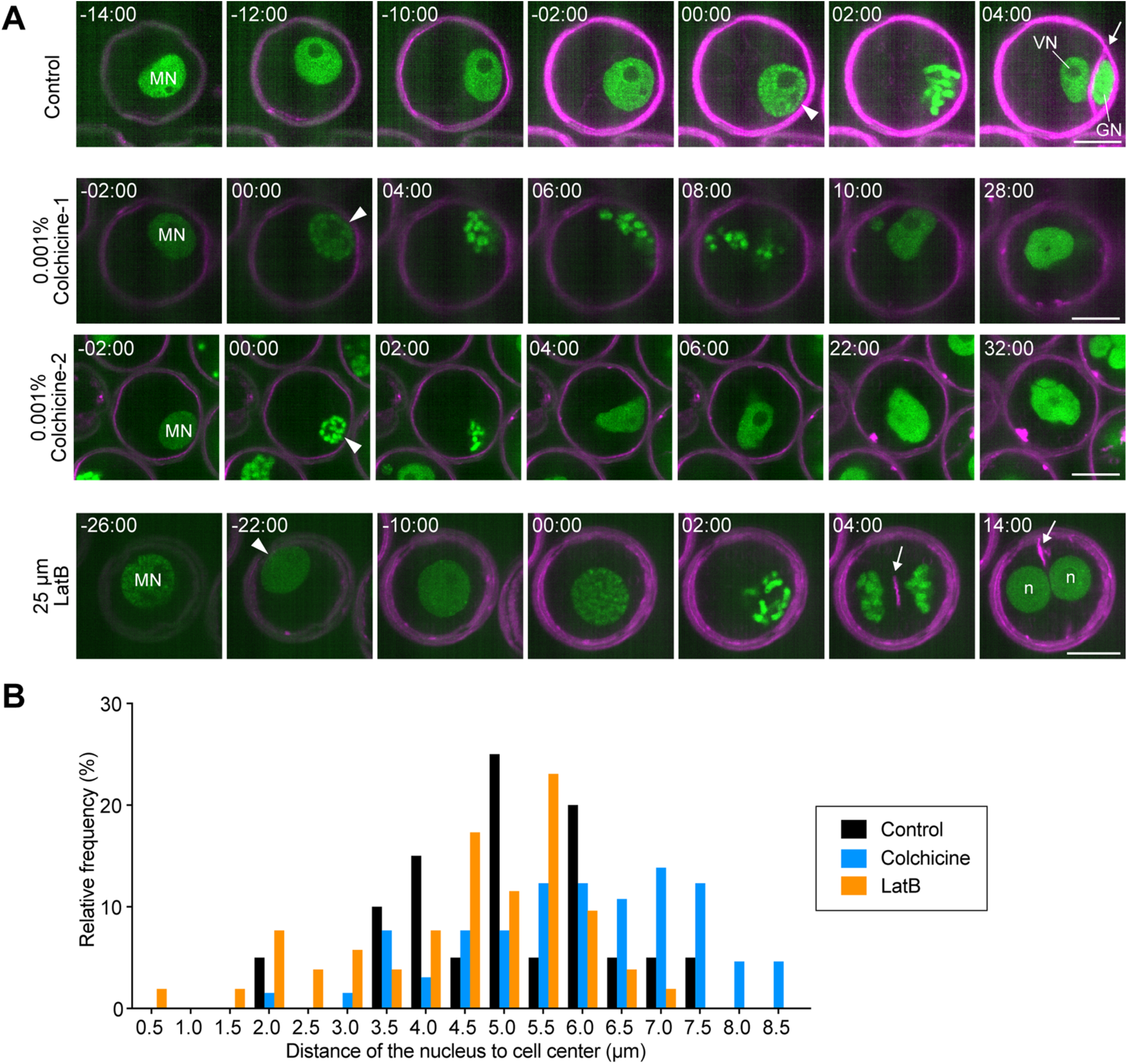
Site and orientation of cell division were defective during PMI under LatB treatment. (A) Confocal live imaging of the nuclei and plasma membrane during PMI with the control, 0.001% colchicine, and 25 μM LatB. The nuclei and plasma membranes were labeled with *AtUBQ10p::H2B-mClover* (green) and PlasMem Bright Red (magenta), respectively. The time when the prophase began was defined as 0 min. White arrows indicate the cell plates during cytokinesis. VN, vegetative cell nucleus; G, generative nucleus. Scale bars, 10 μm (A). (B) Distribution of the distance of the cell division site from the cell center at metaphase [*n* = 20 (Control), 65 (Colchicine), and 52 (LatB)]. The cell division sites were calculated from the position of the chromosome at metaphase. The cell center was calculated using the signals from PlasMem Bright Red.

### After asymmetric division, the position and size of the vegetative nucleus changes dynamically, whereas that of the generative nucleus remains almost the same

To analyze nuclear dynamics after PMI, confocal live imaging was performed using microspores from the nuclear marker line (Figure 4A). Images were captured every 30 min from before PMI to 48 h after PMI, and the resulting z-stack images were projected to track the nuclear positions and measure the sizes of vegetative and generative nuclei in bicellular pollen (Movie 5). Because the pollen was spherical, only microspores with an axis located in the *xy*-plane were used to analyze the images. The size of the microspore nuclei during the G2 phase immediately before PMI was defined as 1.0, and the nuclear areas before and after PMI were measured (Figure 4B). During anaphase, the areas of sister chromatids were approximately 0.25 and then increased rapidly to 0.4–0.6 by the second hour after PMI (Figure 4A and 4B; Control). Thereafter, the size of the vegetative nucleus continued to increase linearly, reaching 1.50 at 40 h, whereas that of the generative nucleus was maintained at approximately 0.5 at 40 h (Figure 4B; Control).

**Figure 4.**
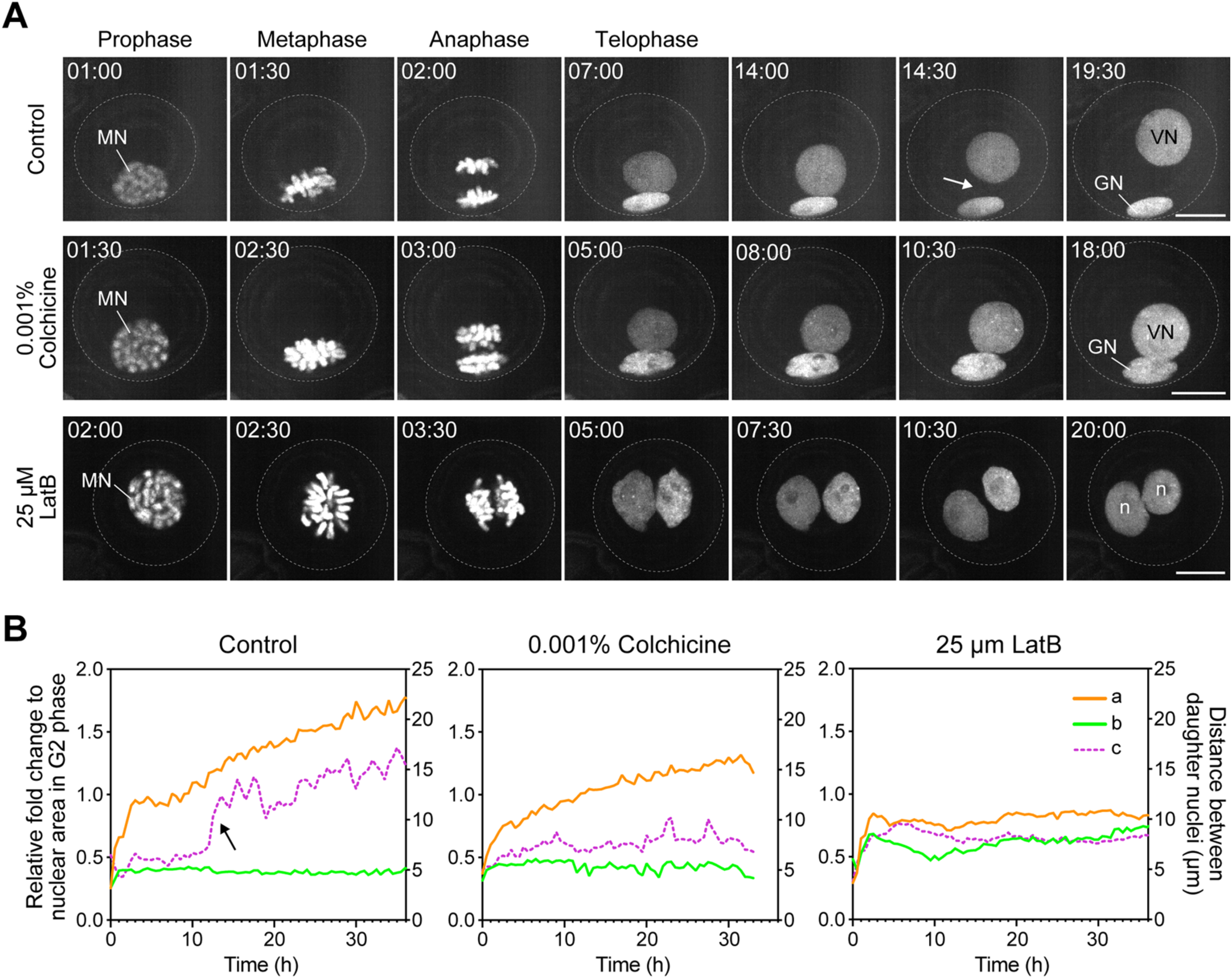
Nuclear dynamics and geometry were affected after PMI with cytoskeletal inhibitors. (A) Confocal live imaging of nuclei dynamics from prophase to G1 phase with the control, 0.001% colchicine, and 25 μM LatB. Nuclei were labeled with *AtUBQ10p::H2B-mClover* (green). The time when the prophase began was defined as 0 min. MN, microspore nucleus; VN, vegetative cell nucleus; GN, generative nucleus; n, daughter nuclei after PMI. Scale bars, 10 μm. (B) Relative area (left; a, b) and distance between daughter nuclei (right; c) after chromosome segregation. The time at which anaphase was initiated was defined as 0 min. The relative fold change of the daughter nuclei area was calculated using the signals of *H2B-mClove*. The nuclear area of the G2 phase (3 h before chromosome segregation) was defined as 1. Nuclei distant from the cell center are indicated by a, and nuclei close to the center are indicated by b.

Next, we tracked the nuclei in the pollen before and after PMI. In control microspores, after migrating to the generative pole, the nucleus entered prophase while remaining in place and underwent cytokinesis to become bicellular (Movie 5 and Figure 4A; Control). The position of the generative nucleus remained at the generative pole, and its size was maintained for 36 h (arrows in Figure 4A; Control). By contrast, the vegetative nucleus migrated to the inner of the vegetative cell immediately after cytokinesis for approximately 12 h and then promptly moved toward the vegetative pole (arrows in Figure 4A and 4B; Control, 14:30). The vegetative nucleus was located near the vegetative pole at a distance from the generative cell (Figure 4A; Control, 19:30). The timing of such nuclear migration of the vegetative nucleus varied from early to late but was observed in all cases (16 ± 10 h, *n* = 64, Figure 5A; Control). When metaphase onset was set to 0 h, the distance between both nuclei was approximately 5 μm immediately after cytokinesis until 12.5 h (Figure 4B; Control), whereas it rapidly increased to approximately 10 μm and then gradually increased to 15 μm after 35 h (Figure 4B; Control). The size of the vegetative nucleus did not expand rapidly at 12.5 h after metaphase, indicating that no apparent correlation exists between size expansion and migration of the vegetative nucleus (Figure 4B; Control).

**Figure 5.**
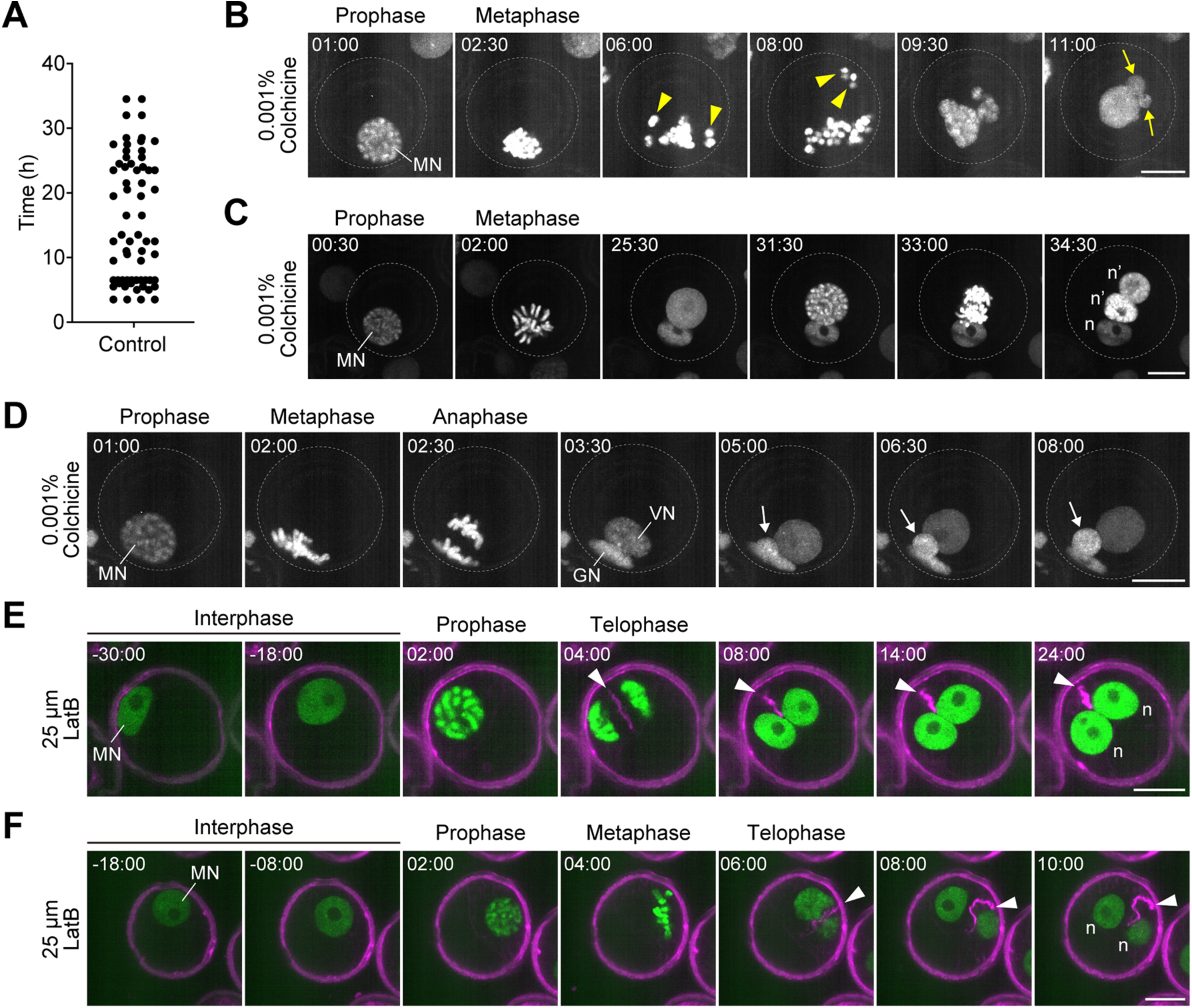
Defects in nuclear dynamics, cell plate formation, and cell cycle progression after cytoskeletal inhibition. (A) Time to rapid nuclear migration of VN from the GN after chromosome segregation [*n* = 64 (Control)]. (B–F) Confocal live imaging of nuclei and plasma membrane from prophase to G1 phase with 0.001% colchicine (B–D) and 25 μM LatB (E, F). The nuclei and plasma membranes were labeled with *AtUBQ10p::H2B-mClover* (green) and PlasMem Bright Red (magenta), respectively. The time when the prophase began was defined as 0 min. Yellow arrowheads and arrows in (B, C) indicate the disassembled chromosomes and micronuclei, respectively. White arrows and arrowheads indicate escape of the generative nucleus and cell plate formation, respectively. MN, microspore nucleus; VN, vegetative nucleus; GN, generative nucleus; n, daughter nuclei after PMI. Scale bars, 10 μm.

### Actin filaments are required for the proper cell plate expansion during cytokinesis and nuclei enlargement after asymmetric division

To examine the effects of microtubules and actin filaments on the regulation of the nuclear position and size, we added each inhibitor and performed observational and image analyses after PMI. When colchicine, a microtubule formation inhibitor, was added, nuclei shifted polarity and asymmetric division were observed in many pollen grains, similar to those in the control (Figure 4A; Colchicine); however, some pollen grains exhibited abnormal chromosome segregation during metaphase (Movie 6 and Figure 5B). These disorganized chromosomes were then reconstructed with nuclear membranes; however, many formed micronuclei instead of binuclei, as observed in the control pollen (arrows in Figure 5B). These micronuclei gradually fused to form a single large nucleus in the pollen. Asymmetric division was also observed in some colchicine-treated microspores, as observed in the control. In these pollen grains, similar to the control, the generative nucleus remained constant at approximately 0.4, whereas the vegetative nucleus gradually increased to approximately 1.25 at 35 h after PMI (Figure 4B; Colchicine). However, unlike the control, the distance between nuclei increased slightly from 5 μm immediately after PMI but remained almost constant at approximately 0.6 within 35 h (Figure 4B). Notably, we also observed that some pollen grains showed vegetative cells entering mitosis and dividing again (Movie 6 and Figure 5C; 31:30, 33:00, and 34:30), and disruption of the cell plate resulted in the escape of the generative nucleus into the vegetative cell (Movie 6 and arrows in Figure 5D) after asymmetric division.

The effects of actin filament disruption after PMI were also investigated, as actin filaments are important in cell plate expansion and guidance in plant cytokinesis (Yi & Goshima, 2022; Zonia et al., 1999). Time-lapse observations revealed that many LatB-treated microspores showed abnormal cytokinesis (Movie 7). Almost all the abnormal pollen grains exhibited nuclear division and subsequent cell plate formation, whereas cell plate expansion was impaired, resulting in the failure of cytokinesis (Figure 5E). The cell plates formed at the center of the division plane did not expand and reached the pollen wall, resulting in cytokinesis failure (arrowheads in Figure 5E). Multinucleate pollen formed two nuclei per cell (Figure 5E; 24:00), and exhibited no successful completion cytokinesis. No pollen was observed that vegetative nucleus underwent an additional division, similar to that observed in the colchicine-treated pollen. The nuclear position and size of pollen grains were measured immediately after chromosome segregation and 35 h later (Movie 5). The distance between both nuclei was approximately 5 μm immediately after cytokinesis until 2.5 h, whereas it rapidly increased to approximately 7–10 μm (Figure 4B; LatB). Unlike in the control and colchicine-treated pollen, the size of the vegetative nucleus did not expand after 35 h. Both daughter nuclei were approximately the same size, approximately 0.6–0.8 (Figure 4B; LatB). Notably, a mismatch between the positions of the division plane and the cell plate was observed in some LatB-treated pollen; that is, the cell plate was formed in a direction perpendicular to the division plane during telophase (arrows in Figure 5F). These results suggest that actin filaments are important for maintaining the cell axis in the microspore, nuclear position, and cytokinesis progression during PMI.

## Discussion

PMI is a simple process that couples cell division and fate determination through asymmetric mitotic cell division and is an excellent example of cellular differentiation from a single cell in plants. Time-lapse imaging revealed that the cell cycle of PMI was 169 min. In *Arabidopsis*, meristemoids are produced by the asymmetric division of the epidermis and then differentiate into guard mother cells, which divide symmetrically to create paired stomatal guard cells (Zhang et al., 2023). The durations of asymmetric division and symmetric division of stomatal cells are 12 and 20.27 h, respectively (Han et al., 2022). By contrast, cell cycle progression in *Arabidopsis* root cells requires 69 min (Pietra et al., 2013). These differences indicate that the duration of the cell cycle depends on the cell type and events, even within the same cell lineage. During pollen development, symmetric division of the generative cell after PMI forms two sperm nuclei. Although the duration of PMII is unclear, differences may exist in cell cycle regulation during pollen development, depending on whether daughter cells differentiate into the same or different cell fates.

Cytokinesis is critical for the architecture of tissues and organs in higher plants because cell division enables the morphogenesis and differentiation of different tissues (Chebli et al., 2021). Our results indicated that both microtubules and actin filaments play fundamental roles in PMI. In land plants, the future division plane of mitotic cells is determined by the cortical ring of microtubules and F-actin, known as the preprophase band (PPB) (Walker et al., 2007). However, PPBs, including micropores, are absent in some mitotic cells (Terasaka & Niitsu, 1987). The mechanism by which the positions of the nucleus and division plane are determined in microspores without PPB remains unknown. Our pharmacological studies indicated that nuclear migration is not disrupted before PMI but is impaired during the anaphase-to-telophase transition in the presence of colchicine. Microtubules radiating from the nuclear membranes have also been reported in microspores from *Gasteria verrucosa* (Van Lammeren et al., 1985). Pharmacological studies using colcemid in *Tulipa gesneriana* and colchicine in *Pinus* microspores suggested that microtubules are not required for nuclear migration but are necessary for nuclear position maintenance (Tanaka & Ito, 1981; Terasaka & Niitsu, 1987). In tobacco pollen, microtubules visualized using TUA6 localize near the nuclear membrane as a basket-like nuclear cap and at the spindle (Oh et al., 2009). The basket-like nuclear cap of microtubules visualized with TUB6 in *N. benthamiana* may maintain the nuclear position prior to division (Figure 2C). After the anaphase, actin filaments are necessary for the primary expansion of the cell plate in tobacco ‘Bright Yellow-2’ (BY-2) cells (van Oostende-Triplet et al., 2017). In our study, microtubule inhibition resulted in the escape of the generative nucleus into vegetative cells (Figure 5D), indicating an impairment in the initially formed cell plate, similar to that observed in BY-2 cells. Furthermore, actin polymerization is required for the centrifugal growth of the cell plate and fusion with the parental plasma membrane (Hepler et al., 2002; van Oostende-Triplet et al., 2017), which is also supported by our observations (Figure 5E and 5F). The 2,3,5-triiodobenzonic acid (TIBA), an inhibitor of actin depolymerization, induces oblique cell plate formation in BY-2 and *Arabidopsis* root cells, indicating that actin filaments are critical in spindle orientation and cell plate formation (Kojo et al., 2013). A mismatch was observed between the directions of the division plane and the cell plate in LatB-treated pollen (Figure 5F). These results suggest that the disruption of actin filaments prevents the maintenance of nuclear position; therefore, the spindle is formed in an incorrect orientation, but the cell plate is formed according to the original axis, which can result in such a mismatch. In addition to cytoskeletal elements, organelles such as mitochondria, vesicles, and vacuoles are important in pollen development (Nagata, 2003). Vacuole functions vary depending on the developmental stage of the pollen and their morphology notably changes during microgametogenesis (Pacini et al., 2011). Further imaging analysis and inhibition of related genes using transient genome editing of microspores (Nagahara et al., 2021) will be useful for understanding the cytoskeletal regulation of microspore development and differentiation.

After cytokinesis, the sizes of the vegetative and generative nuclei were substantially different because the vegetative chromatin was decondensed, whereas the generative chromatin was highly condensed (Figure 4). When actin polymerization was impaired, two daughter nuclei of approximately the same size formed after PMI, resulting in multinucleate pollen (Figure 4B). This indicates that the vegetative nucleus cannot enlarge until the cytoplasmic division is complete. Notably, we observed that the vegetative nucleus of pollen treated with colchicine divided again after PMI (Figure 5C). This suggests that when cytokinesis is not completed normally after PMI and the nucleus fails to gain vegetative cell identity, it retains the capacity of microspores for division. CenH3 is removed from the vegetative cell centromere after PMI to terminally differentiate and exit the cell cycle, whereas it remains in the generative cell centromere to enter PMII (Borg & Berger, 2015). A recent study based on imaging and RNA-seq analysis of *Arabidopsis* sperm cells showed that transcriptionally active regions are highly restricted in sperm chromatin compared to those in vegetative chromatin (Shibuta & Matsunaga, 2023). Additionally, RNA-seq analysis of *Arabidopsis* sperm cells revealed that the promoters of genes that are upregulated in the sperm during pollen tube growth are already accessible in the sperm chromatin of mature pollen grains (Misra et al., 2025). These results indicate that chromatin condensation and decondensation are important for cell-specific gene expression and that the regulation of chromatin structure is responsible for cell cycle termination/progression and differentiation of both daughter cells after PMI. Transcription from the haploid genome is considerably activated during PMI, leading to a transition from the sporophyte to gametophyte phases (Nelms & Walbot, 2022). Further comprehensive studies on cell morphology, including the cytoskeleton, gene expression, protein function, and nuclear chromatin structures, during PMI will reveal the regulatory mechanisms of asymmetric division and differentiation in pollen development.

## Materials and methods

### Plant materials

In this study, we used wild-type and transgenic *N. benthamiana AtUBQ10p:: H2B-mClover*, as previously described (Nagahara et al., 2021). The growth conditions have been described previously (Kaneshiro et al., 2022). To prepare microspores and immature pollen grains, anthers were manually isolated from each bud before anthesis at stages 1 and 2, which were defined as the early and late stages, respectively (Steinbachová et al., 2021).

### Pollen culture

For live imaging, a pollen culture medium was used according to a previous study (Tupý et al., 1991): 0.5 M sucrose, 3 mM glutamine, 10 g/L lactalbumin hydrolysate, 10 mM KNO_3_, 1 mM Ca (NO_3_)_2_·4H_2_O, 1 mM MgSO_4_·7 H_2_O, 0.16 mM H_3_BO_3_, 1 mM uridine, 0.5 mM cytidine, and 1 mM phosphate buffer (pH 7.0). Following sterilization using a syringe filter (Millex-GV 0.22 µm PVDF; Merck, Darmstadt, Germany), the culture medium was stored at −30 °C until use. Pollen grains were released by gently squashing the anthers using sterilized tweezers in a pollen culture medium on a clean bench. The pollen culture medium was poured into a multi-well glass-bottom dish (D141400; Matsunami Glass, Osaka, Japan).

### Plasmid vectors used for the transient expression

The fluorescent nuclear reporter plasmids, *35Sp:H2B-tdTomato* (YMv199) and *AtUBQ10p:H2B-tdTomato* (YMv266), have been previously described (Kaneshiro et al., 2022). Plasmid vectors, that have been previously reported for *pUBQ10:EYFP-TUB6,* were used to label the microtubules (Sasaki et al., 2023). Plasmid vectors that have been previously reported for *UBQ10pro:LifeAct-mGFP* were used to label the actin filaments (Kijima et al., 2025).

### Biolistic delivery of plasmid DNA into pollen

Anthers from the bud at stage 1 (Steinbachová et al., 2021) before anthesis were collected using tweezers immediately before bombardment. Three fresh anthers were squashed onto a cut wet hydrophilic PTFE membrane (approximately 2.2 cm^2^; JCWP04700 Omnipore, MERCK, Darmstadt, Germany) with 20 μL pollen culture medium to spread the microspores (Kaneshiro et al., 2022). Biolistic delivery of plasmid DNA was performed using the PDS-1000/He system (Bio-Rad Laboratories, Hercules, California, USA) according to a previously described method (Nagahara et al., 2021). To visualize bombarded pollen, a maximum of 1 μg of DNA containing a single plasmid or a mixture of 500 ng of each plasmid was used. After bombardment, 200 μL of pollen culture medium was added to the filter to suspend pollen. The pollen suspension was then equally divided and transferred into two wells of a four-well 35-mm glass base dish (D141400; Matsunami Glass Ind., Ltd., Japan), sealed with Parafilm (Bemis Flexible Packaging, Oshkosh, WI, USA), and cultured at 25–30 °C in an incubator as previously described (Kaneshiro et al., 2022). For incubations longer than 20 h, sterilized equipment was used in all the above processes, which were performed on a clean bench. Plasmid-introduced pollen was defined based on the expression of fluorescent proteins that could be detected after at least 4 h, as previously described (Nagahara et al., 2021).

### Inhibitor

The microspore population from the nuclear marker *AtUBQ10p::H2B-mClover* was used for pharmacological studies on developmental progression during PMI. To disrupt microtubules, microspore suspensions were treated with 0.001% (w/v) colchicine (AdipoGen Life Sciences, Inc., CA, USA) for 5 min. Similarly, to disrupt actin filaments, microspores were treated with 25 μM LatB (AdipoGen Life Sciences, Inc., CA, USA).

### Cell membrane staining

To visualize the cell membranes, PlasMem Bright Red solution (Dojindo Laboratories, Kumamoto, Japan) was added to the culture medium at a final concentration of 0.15% (v/v). Microspores were suspended in a culture medium containing PlasMem Bright Red, poured into a multi-well grass-bottom dish (D141400; Matsunami), and incubated for more than 2 min. After incubation, the culture dishes were observed under a microscope for time-lapse imaging.

### Imaging analysis to assess pollen viability

To analyze the pollen viability, an inverted fluorescent microscope (Eclipse Ti2-E, Nikon, Tokyo, Japan) equipped with an objective lens (CFI Plan Apo λ 20×, NA = 0.75, WD = 1.0 mm; CFI Plan Fluor 40×, NA = 0.75, WD = 0.66 mm) was used. The emitted fluorescence signals were detected using a CMOS camera (ORCA-Fusion BT C15440-20UP; Hamamatsu Photonics, Shizuoka, Japan). Bandpass filters, including fluorescein isothiocyanate (FITC) for the mClover signal and Cy3 for tdTomato fluorescence (Nikon), were used to capture the images. To assess pollen viability, microspores from *AtUBQ10p:: H2B-mClover* were suspended in a culture medium and observed immediately. The images were analyzed using NIS-Elements AR version 5.42 (Nikon). Based on the fluorescent signals, pollen with nuclei labeled with mClover was defined as viable pollen, whereas red autofluorescence with a shriveled shape was defined as aborted pollen. Among the living pollen grains, 10 pollen grains that showed PMI were selected and the mean diameters were analyzed using Fiji software (Schindelin et al., 2012).

### Confocal live imaging of pollen development

Confocal images were acquired using an Eclipse Ti2-E microscope (Nikon) equipped with a spinning disk confocal scanning unit (CSU-W1; Yokogawa, Tokyo, Japan), 488 and 561 nm lasers (YHQ-LC-488-50-1; YHQ-LC-561-50-1; Yokogawa Electric Co.), an objective lens (CFI SR HP Plan Apo Lambda S 100×C Sil, NA = 1.35, WD = 0.31–0.29 mm; Nikon), and a scientific CMOS camera (ORCA-Fusion BT C15440-20UP; Hamamatsu Photonics). For time-lapse imaging, sequential images were acquired using 488 and 561 nm lasers every 15–120 min. Time-lapse images were processed using NIS-Elements AR and analyzed using Fiji software.

### Imaging analysis of the nuclear tracking

We used the object-tracking function in the NIS-Elements AR software with binary images to track the nuclear position, area, and cell center of the pollen. To create binary images of H2B-mClover and PlasMem Bright Red, automatic recognition was performed using the Segment Objects.ai function in NIS-Elements AR software.

### Data analysis

Data were manipulated using the Python Pandas package (The pandas development team, 2024). The plots were generated using GraphPad Prism, version 10 (GraphPad Software, San Diego, CA, USA).

## Acknowledgements

We thank Drs. T. Higashiyama, S. Nagahara, S. Yamaoka, D. Maruyama, K. Ebine, and H. Takeuchi for helpful suggestions and discussion. Microscopy was performed at the WPI-ITbM of Nagoya University and supported by the Advanced Bioimaging Support in MEXT/JSPS KAKENHI (22H04926). This work was supported by the Japan Society for the Promotion of Science [Grant-in-Aid for Transformative Research Areas (20H05778 and 20H05779 to Y. M.), Grant-in-Aid for Scientific Research (B) (24K02031 to Y. M.), Asahi Glass Foundation (to Y. M.), and the Japan Science and Technology Agency [FOREST program (JPMJFR233V to Y. M., JPMJFR204T to D.K.)].

## Supplementary material legends

**Movie 1.** *In vitro* live imaging of pollen mitosis I (PMI) in *Nicotiana benthamiana*. Nuclei labeled with *AtUBQ10p::H2B-mClover* are shown in green. Pollen autofluorescence is shown in magenta. The numbers stamped in each image indicate the time (hh:mm) from the beginning of the observation. Scale bar, 20 μm. See Figure 1B for additional details.

**Movie 2.** Confocal live imaging of microtubule dynamics during PMI by the transient introduction of the plasmid markers *AtUBQ10p:H2B-tdTomato* and *pUBQ10:EYFP-TUB6*. The *xy*-maximum projection images in 1-μm steps with two planes are shown. The numbers stamped in each image indicate the time (hh:mm) from the beginning of prophase. Scale bar, 10 μm. See Figure 2C for additional details.

**Movie 3.** Confocal live imaging of actin filaments during PMI by the transient introduction of the plasmid markers *AtUBQ10p:H2B-tdTomato* and *UBQ10pro:LifeAct-mGFP*. The *xy*-maximum projection images in 1-μm steps with two planes are shown. The numbers stamped in each image indicate the time (hh:mm) from the beginning of prophase. Scale bar, 10 μm. See Figure 2D for additional details.

**Movie 4.** Confocal live imaging of the nuclei and plasma membrane during PMI with the control, 0.001% colchicine, and 25 μM LatB. Nuclei and plasma membranes were labeled with *AtUBQ10p::H2B-mClover* (green) and PlasMem Bright Red (magenta), respectively. The numbers stamped in each image indicate the time (hh:mm) from the beginning of prophase. Scale bars, 10 μm. See Figure 3A for additional details.

**Movie 5.** Image analysis of the relative area and distance between daughter nuclei after chromosome segregation with the control, 0.001% colchicine, and 25 μM LatB. Automatically defined areas of the nuclei of the microspore, generative cell, and vegetative cell are colored overlays with blue, green, and orange, respectively. The trajectory of the center of gravity of each nucleus is shown as a solid line with a rainbow color, indicating the movement speed. The numbers stamped in each image indicate the time (hh:mm) from the beginning of prophase. Scale bars, 10 μm. See Figure 4 for additional details.

**Movie 6.** Confocal live imaging of nuclei in the two samples from prophase to G1 phase with 0.001% colchicine. Nuclei were labeled with *AtUBQ10p::H2B-mClover* (green). The numbers stamped in each image indicate the time (hh:mm) from the beginning of prophase. Scale bars, 10 μm. See Figure 5B−D for additional details.

**Movie 7.** Confocal live imaging of nuclei in the three samples from prophase to G1 phase with 25 μM LatB. Nuclei and plasma membranes were labeled with *AtUBQ10p::H2B-mClover* (green) and PlasMem Bright Red (magenta), respectively. The numbers stamped in each image indicate the time (hh:mm) from the beginning of prophase. Scale bars, 10 μm. See Figure 5E and 5F for additional details.

## Notes

### Competing Interest Statement

The authors have declared no competing interest.

